# Mapping First *In Vivo* Ebolavirus Infection Events

**DOI:** 10.64898/2026.09.16.752249

**Authors:** Jiani Liu, Brady N. Zell, Lin Wang, Michael A. Barry

## Abstract

Orthoebolaviruses, including Ebola virus (EBOV) and Bundibugyo virus (BDBV), cause sporadic outbreaks and devastating hemorrhagic infections in humans. A significant challenge that impedes the study of filoviruses is that the infectious virus can only be handled in biosafety level 4 (BSL4) facilities due to their extreme virulence. Life cycle modeling systems that use minigenomes and transcription/replication-competent virus-like particles (trVLPs) are powerful tools for studying these virus’ biology and to explore antiviral therapies under BSL2 conditions. To date, these powerful tools have only been utilized in vitro in cell culture systems. In this work, we demonstrate the use of EBOV trVLPs in vivo. We report the fine specificity of first infection events in a model of EBOV injection in the skin. Cre recombinase expressing monocistronic EBOV trVLPs were packaged and used to inject Cre reporter mice intradermally (ID). Initial infection events were monitored in living animals by luciferase imaging and then by flow cytometry to track cell-specific expression of Cre-activated GFP. By this approach, EBOV trVLPs were shown to directly infect T cells, monocytes, dendritic cells (DCs) with notably high first infections of skin-resident Langerhans cells (LCs) in the skin barrier and its draining lymph node. EBOV trVLPs were next tested as an *in vivo* tool to assess antiviral therapies by testing the ability of an EBOV vaccine to reduce trVLP luciferase levels after infection in the skin. This work demonstrates first *in vivo* use of *EBOV* trVLPs and provides proof of concept for their use as an *in vivo* BSL2 model to more rapidly develop antiviral agents and vaccines.

## Introduction

Orthoebolaviruses including Ebola virus (EBOV) and Bundibugyo virus (BDBV) cause severe hemorrhagic fever with case-fatality rates that can exceed 50% [1–5]. Understanding viral dissemination, tissue tropism, and host immune responses is critical for developing effective vaccines and antiviral therapies. However, studies involving infectious EBOV require biosafety level 4 (BSL4) containment, greatly limiting mechanistic investigations and preclinical evaluation to the few BSL4 research laboratories. This high biosafety requirement limits not only basic research into these orthoebolavirus*’* biologies but drastically impedes screening and evaluating antiviral and immunologic countermeasures against these devastating pathogens.

To overcome these limitations, several surrogate life cycle modeling systems have been developed [6–8] in which transcription- and replication-competent virus-like particles (trVLPs) faithfully recapitulate multiple steps of the EBOV replication cycle while remaining compatible with BSL2 laboratories [7, 9]. *Orthoebolavirus* trVLPs have become indispensable *in vitro* tools for investigating viral transcription and replication strategies, as well as host-pathogen interactions, and antiviral interventions. While trVLPs are useful for cell culture studies, they have not been utilized for *in vivo* life cycle modeling or to evaluate antiviral drugs or vaccines against orthoebolaviruses.

To address this research gap, we harnessed a sensitive *in vivo* reporter gene platform that we previously developed to fingerprint gene therapy vector tropism in mouse models [10–13]. In this system, vectors expressing Cre recombinase are delivered into “mT/mG” mice and/or “LSL-Luc” mice (**Fig. 1**).

**Figure 1.**
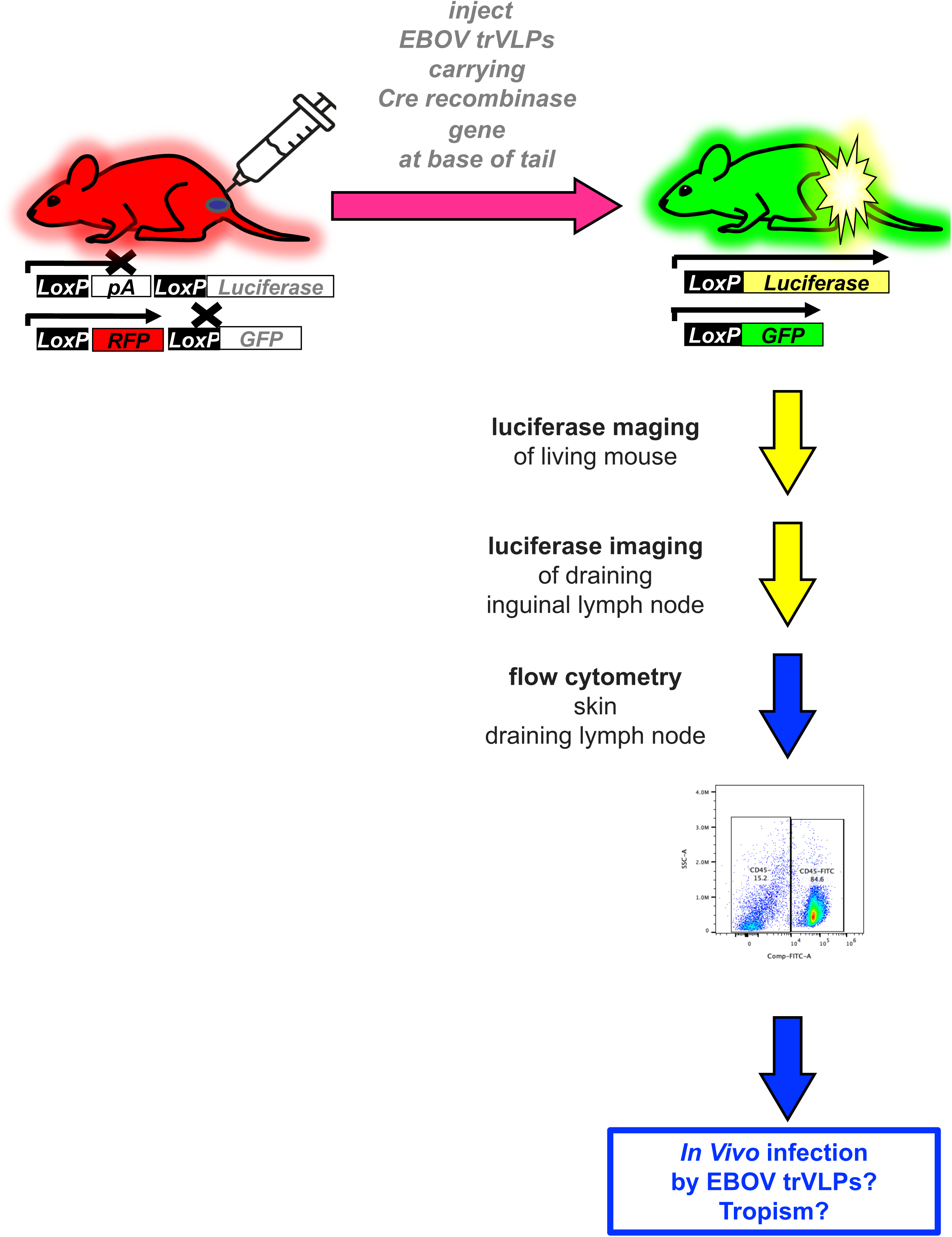
Testing of the Cre Reporter System for *In Vivo* Tracking of EBOV trVLPs. Cre-reporter mice carrying Cre-inducible luciferase and GFP reporters were injected intradermally at the base of the tail with EBOV trVLPs carrying Cre recombinase. Cre-mediated recombination activates reporter expression, enabling detection by luciferase bioluminescence imaging and GFP fluorescence. Luciferase activity was monitored longitudinally by in vivo imaging, followed by imaging of the draining inguinal lymph nodes. Skin from the injection site and draining inguinal lymph nodes were subsequently collected and processed for flow cytometric analysis of GFP-positive cell populations to characterize cellular tropism.

In LSL-Luc mice, a luciferase transgsene in the Rosa26 locus is inactivated by an upstream LoxP-poly adenylation-LoxP sequence. If Cre is delivered into the mouse’s cells, Cre deletes this “floxed” cassette allowing transcription and expression of luciferase for *in vivo* imaging of living animals, *ex vivo* imaging of their organs, or highly sensitive enzymatic *in vitro* luciferase assays [10–14].

mT/mG are transgenic at the same Rosa26 locus for a loxP flanked (floxed) membrane-targeted Tomato (mT) red fluorescence protein open reading frame (ORF) that is followed by the ORF for membrane-targeted GFP (mG). In the absence of Cre, mT is expressed and mG is not. Without Cre, the membranes of all of the cells in the mice are red fluorescent. If Cre is delivered into the cell by a vector, mT is deleted and mG is activated turning the cell’s membrane from red fluorescence to green fluorescence. GFP activation can be monitored by IVIS imaging [13], but it is more powerfully used to scrutinize the identity of cells that received Cre via vector delivery using confocal microscopy and flow cytometry. Comparison of mT and mG membrane fluorescent provides exquisite separation and identification of vector modified cells to the extent that in some cases, red membranes of one cell can be seen abutted against the green membranes of an adjacent cell [10].

When mT/mG and LSL-Luc mice are crossed, their progeny manifest both Cre activation phenotypes allowing *in vivo* imaging as well as precise cell identification by confocal microscopy and flow cytometry by non-viral and viral vectors, mRNA, DNA, adenovirus, and adeno-associated virus vectors [10–14]. Subsequent work by another group showed that a Cre vector in this type of system is 8-fold more sensitive than directly delivery of GFP-expressing vectors in mice [15].

Given the utility of the Cre reporter system to track adenovirus, adeno-associated virus, and lipid nanoparticles *in vivo,* we tested its utility to study EBOV trVLPs in this work (**Fig. 1**). To do this, we tested the *in vivo* tropism of EBOV Cre trVLPs in mT/mG and LSL-Luc mice in a model of cutaneous infection. Following this, we tested if EBOV Cre trVLPs could be used as a challenge virus to test the efficacy of an EBOV vaccine.

## Results

### *In Vitro* Production of EBOV Cre trVLPs

To validate the Cre-dependent reporter system *in vitro*, Cre-activated red/green (R/G) reporter 293 cells were used during trVLP production and monitored for fluorescence conversion over time. 293 R/G cells were co-transfected with one antisense EBOV minigenome driven carrying the Cre cDNA that is expressed from a T7 promoter along with 8 other plasmids expressing EBOV helper proteins and T7 polymerase. Following transfection of these passage 0 (P0) cells with the trVLP plasmids, cells in the monolayer were progressively converted from red to green fluorescence over time (**Fig. 2A**). The fraction of GFP+ cells increased over time demonstrating significant time-dependent increases in reporter activation from the EBOV minigenome in the P0 Cre reporter cells (**Fig. 2B**).

**Figure 2.**
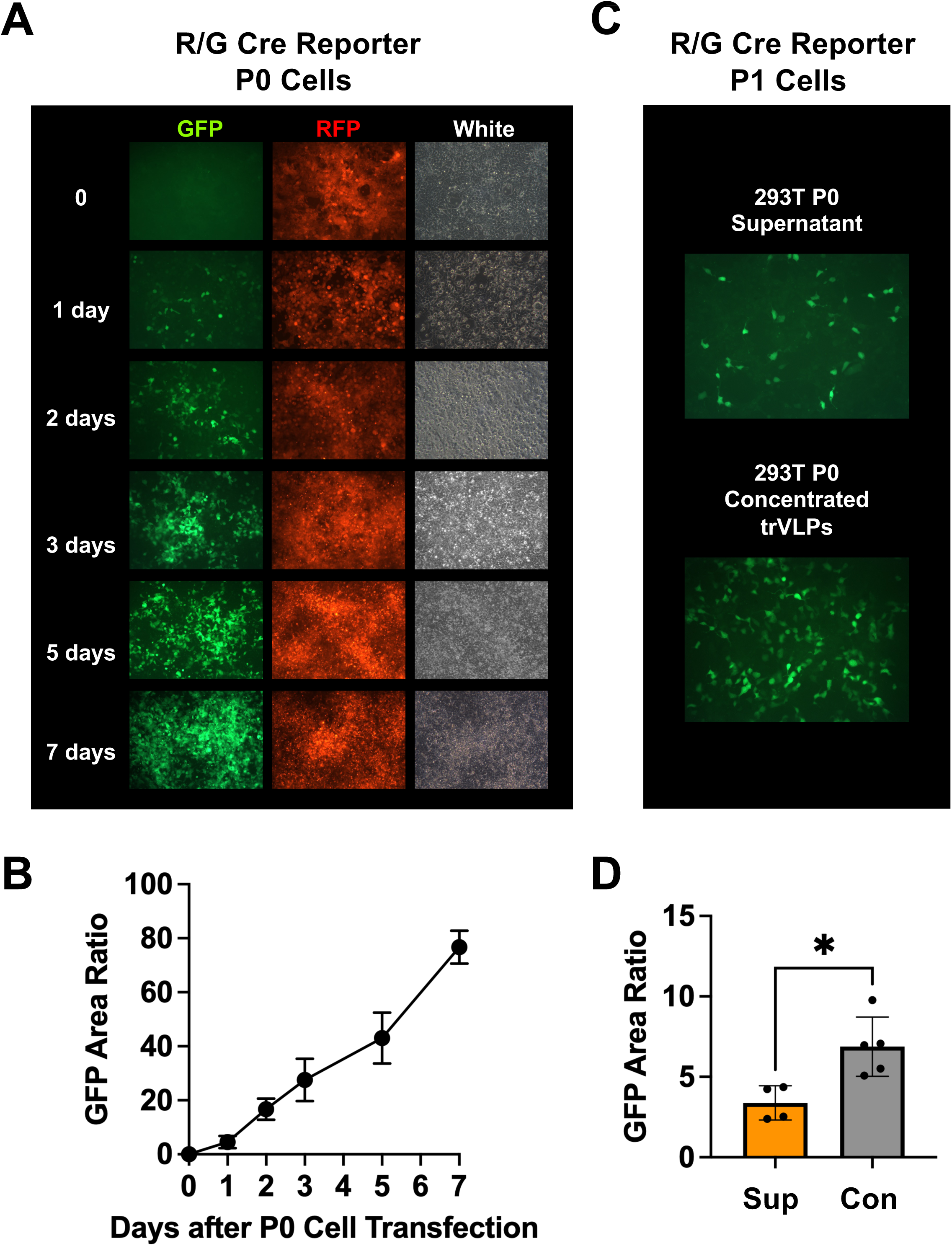
Cre Reporter Activation in Transfected P0 Cells and P1 Reporter Cells by EBOV trVLPs. **A)** Representative fluorescence and bright-field images of R/G Cre-reporter cells during P0 trVLP production. Cells were monitored at the indicated time points (days 0, 1, 2, 3, 5, and 7), showing progressive conversion from red fluorescence to GFP following Cre recombination. GFP, RFP, and corresponding bright-field images are shown. **B)** Quantification of GFP fluorescence intensity in P0 reporter cells over time. Data are presented as mean ± SD from three independent experiments. Statistical comparisons were performed using ordinary one-way ANOVA followed by Dunnett’s multiple-comparisons test, with each time point compared with the control PBS treatment group. By day 7, *p* < 0.0001. **C)** Representative GFP fluorescence images of P1 reporter cells following exposure to P0 culture supernatant or concentrated trVLP preparations. (D) Quantification of GFP fluorescence intensity in P1 reporter cells 72 h after treatment. Statistical significance was determined using a two-tailed unpaired t-test with Welch’s correction. *P* < 0.05 was considered statistically significant.

When cell supernatant or gradient-concentrated Cre trVLPs produced from non-fluorescent 293T P0 cells were applied to 293 R/G Cre reporter cells, GFP+ cells were produced (**Fig. 2C**) with higher numbers of GFP-expressing cells observed in the concentrated trVLPs (p < 0.05 by T test, **Fig. 2D**).

### *In Vivo* EBOV trVLP Activity in a Model of Skin Infection

Although EBOV trVLP systems have been extensively used for *in vitro* studies, to our knowledge, they have not been tested for activity *in vivo*. Given the ability of Cre trVLPs to activate reporter gene in cell culture, we tested whether they could activate luciferase and GFP *in vivo* in mice after intradermal injections as a model of EBOV infection in the skin. It is thought that some EBOV infections occur at breaks in the skin’s barrier [16].

P0 Cre trVLP cell supernatants and sucrose gradient-concentrated trVLPs were injected intradermally (ID) at the base of the tails of mice (**Fig. 4**). This site was chosen, since we previously demonstrated that injection of various gene therapy vectors at this site leads to drainage of vectors and antigen-presenting cells (APCs) to the inguinal lymph node [14].

These ID trVLP injections induced progressive increases in luciferase activity in the skin of the mice over 5 to 7 days by IVIS imaging (**Fig. 3A**). Mice that received unconcentrated trVLP-containing supernatant exhibited lower luciferase activity than those receiving concentrated trVLPs similar to their relative activities *in vitro.* When whole-body total luciferase activities were measured, this revealed significant effects of treatment, time, and treatment × time interaction (**Fig. 3B**). In particular, concentrated trVLPs generated significantly greater reporter activity than PBS on days 3, 5, and 7.

**Figure 3.**
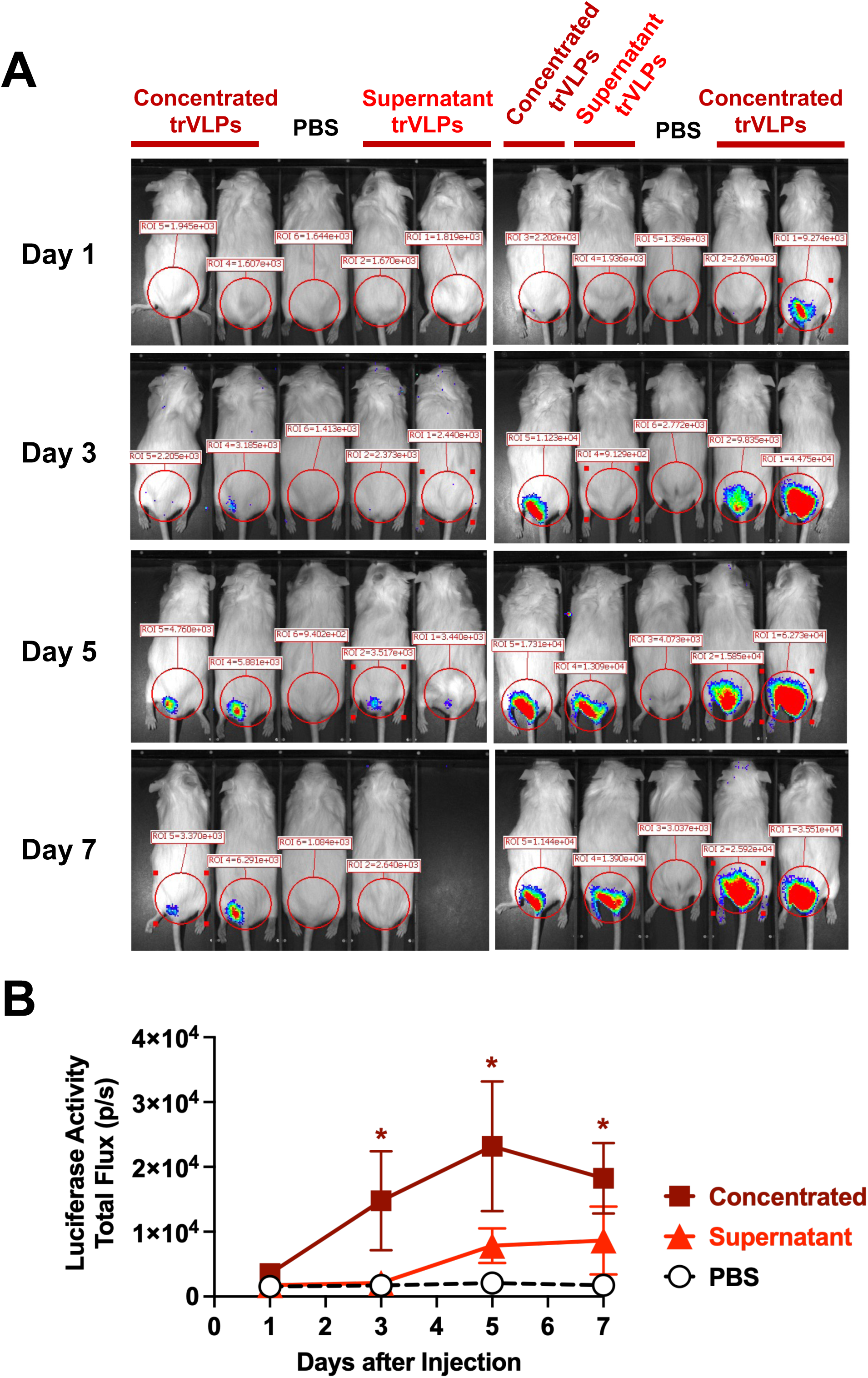
Longitudinal Luciferase Imaging of EBOV trVLP Infections *In Vivo*. **A)** Representative whole-body bioluminescence images of Cre reporter mice following intradermal injection of concentrated trVLPs, unconcentrated trVLP-containing supernatant, or PBS. Mice were imaged on days 1, 3, 5, and 7 after injection. Red circles indicate the regions of interest (ROIs) used for quantification of luciferase activity at the injection site. **B)** Quantification of luciferase activity as total flux within the defined ROIs. Data are presented on the original scale as mean ± SEM. Total flux values were log10-transformed prior to statistical analysis. Data were obtained from two experimental cohorts receiving 100 µg or 200 µg of the respective trVLP preparations. Radiance Color scale min=1.15e^4^ max=4.15e^4^. Longitudinal data were analyzed using a repeated-measures mixed-effects model (REML), followed by Tukey’s multiple-comparisons test for comparisons between treatment groups to the control group at each time point. Significant effects of treatment (*p* = 0.0112), time (*p* = 0.0002), and treatment × time interaction (*p* = 0.0037) were observed. Concentrated trVLPs produced significantly greater luciferase activity than PBS on days 3, 5, and 7. * *p* < 0.05 versus PBS at the corresponding time point.

At the end of the study, the animals were euthanized and their inguinal draining lymph nodes were imaged for *in situ* luciferase activity (**Supplemental Fig. 1A**). Luciferase activity was detectable in these draining lymph nodes in a subset of trVLP-treated animals. However, variations between the animals did not reveal statistically significant difference among treatment groups (**Supplemental Fig. 1B**).

### *In Vivo* Cell Tropism of EBOV trVLPs in the Skin

The skin and inguinal draining lymph nodes were collected after euthanasia and their cells analyzed by multiparameter flow cytometry. Representative gating strategies are shown in **Supplemental Fig. 2 and 3**. After the exclusion of debris, cell doublets, and nonviable cells, leukocyte populations were identified using a combination of antibodies against CD45, CD11b, CD11c, MHC-II, CD207, and CD3. These cells were examined to determine if they were GFP-positive or GFP-negative to assess the cell tropism of the trVLPs in the skin and its draining lymph node.

Low but detectable GFP activation was observed after injection of unconcentrated trVLPs in the skin (**Fig. 4A**). Higher and significant GFP was observed in immune cells in the skin after exposure to the concentrated trVLPs. CD45⁻ non-immune cells were largely GFP negative, whereas CD45+ immune cells in the skin were GFP-positive. When specific lineages of immune cells were examined, reporter-positive cells were detected among CD45⁺CD3⁺ T cells, CD45⁺CD11b⁺ monocyte/macrophage cells, CD207⁺ Langerhans cells (LCs), CD45⁺CD11c⁺MHC-II⁺ dendritic cells (DCs), CD45⁺CD11c⁺MHC-II⁺CD207⁺ and Langerhans dendritic cells (**Fig. 5A**). Quantification across individual animals confirmed increased GFP positivity in several of these immune-cell populations after trVLP exposure, with highest reporter protein levels generally observed in mice receiving concentrated trVLPs.

**Figure 4.**
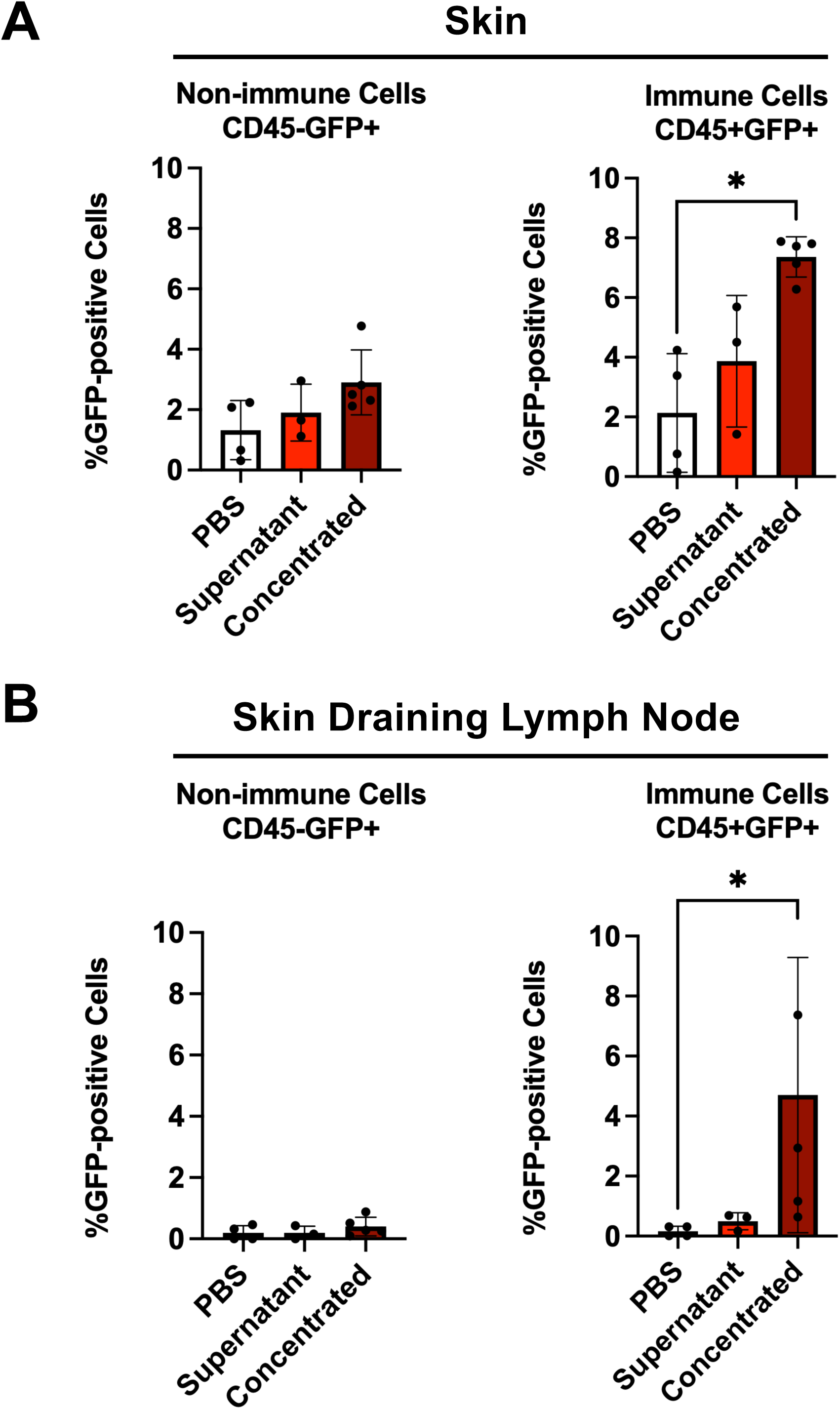
GFP Reporter Activation in Immune and Non-immune Cells Following EBOV trVLP Administration. Quantification of GFP-positive CD45⁺ immune cells and CD45⁻ non-immune cells in **A**) skin and **B**) draining inguinal lymph nodes. Each point represents an individual mouse. Data are presented as mean ± SD. Statistical comparisons among groups were performed using the Kruskal–Wallis test followed by Dunn’s multiple-comparisons test. *p* < 0.05 was considered statistically significant; non-significant comparisons are not indicated.

**Figure 5.**
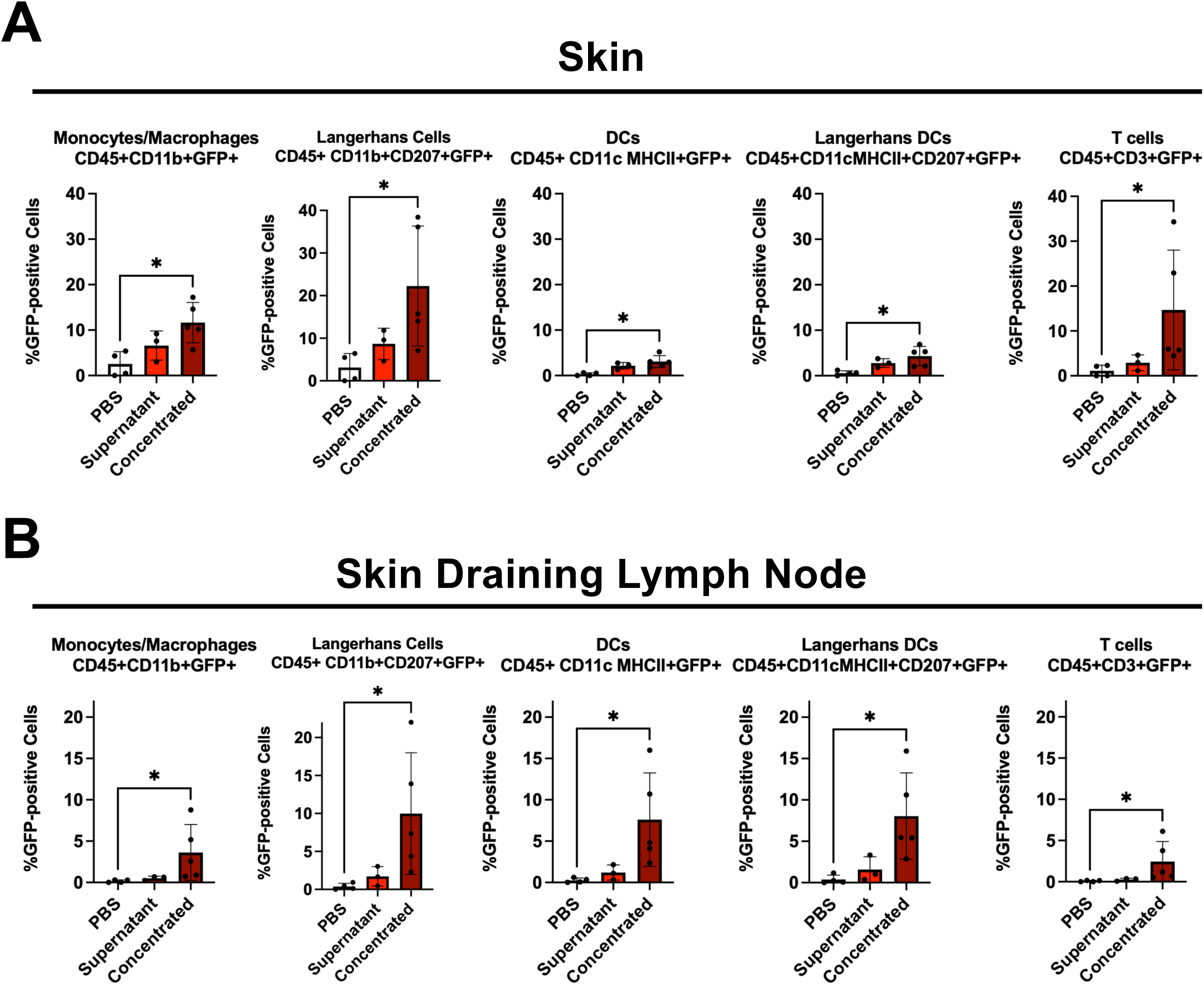
GFP Activation in Immune-cell Subsets by Cre trVLPs. Quantification of GFP-positive cells within the indicated immune-cell subsets, including CD11b⁺ monocytes/macrophages, CD207⁺ Langerhans cells, CD11c⁺MHC-II⁺ dendritic cells, CD11c⁺MHC-II⁺CD207⁺ Langerhans dendritic cells, and CD3⁺ T cells, in **A)** skin and **B)** draining inguinal lymph nodes. Each point represents an individual mouse. Data are presented as mean ± SD. Statistical analysis was performed as described for Figure 4.

### *In Vivo* Cell Tropism of EBOV trVLPs in Skin Draining Lymph Nodes

A similar pattern of infection was observed in cells from the inguinal lymph nodes that drained from the ID injection site (**Fig. 4B and 5B**). GFP-positive cells were again identified within CD45⁺ leukocytes, including CD11b⁺ monocyte/macrophage-associated cells, CD207⁺ LCs, CD11c⁺MHC-II⁺ DCs, CD11c⁺MHC-II⁺CD207⁺ LC/DC cells, and CD3⁺ T cells. In contrast, CD45⁻ non-immune cells showed less GFP than the CD45+ immune cells. Quantitative analysis demonstrated higher GFP positivity across several immune-cell subsets in trVLP-treated animals, particularly following administration of concentrated trVLPs.

### Effects of Ebola Glycoprotein and Polymerase on EBOV trVLPs

Ebola trVLPs were next produced by transfection with all helper protein plasmids with or without the glycoprotein (GP) or polymerase (L) plasmids and supernatants from these cells were used to infect P1 293 R/G cells (**Fig. 6A**). The trVLPs produced from 293T or 293 R/G cells mediated detectable GFP expression in P1 293 R/G cells. In contrast, supernatants from transfections lacking either GP or L produced no GFP in P1 cells. These supernatants were concentrated on sucrose gradients and subjected to silver staining and western blotting. Silver staining for total proteins detected multiple bands in the concentrated samples (**Fig. 6B and Supplemental Figure 5**). Western blotting of the same concentrated samples detected VP24, VP35, VP40, and NP in all of the samples (**Fig. 6C and Supplemental Figure 6**). Notably, glycoprotein was detected in all of the concentrated trVLP samples with the exception of those that were produced without the GP helper plasmid. In contrast, staining for L was equivocal for all samples, consistent with the general difficulty in detecting this large EBOV protein.

**Figure 6.**
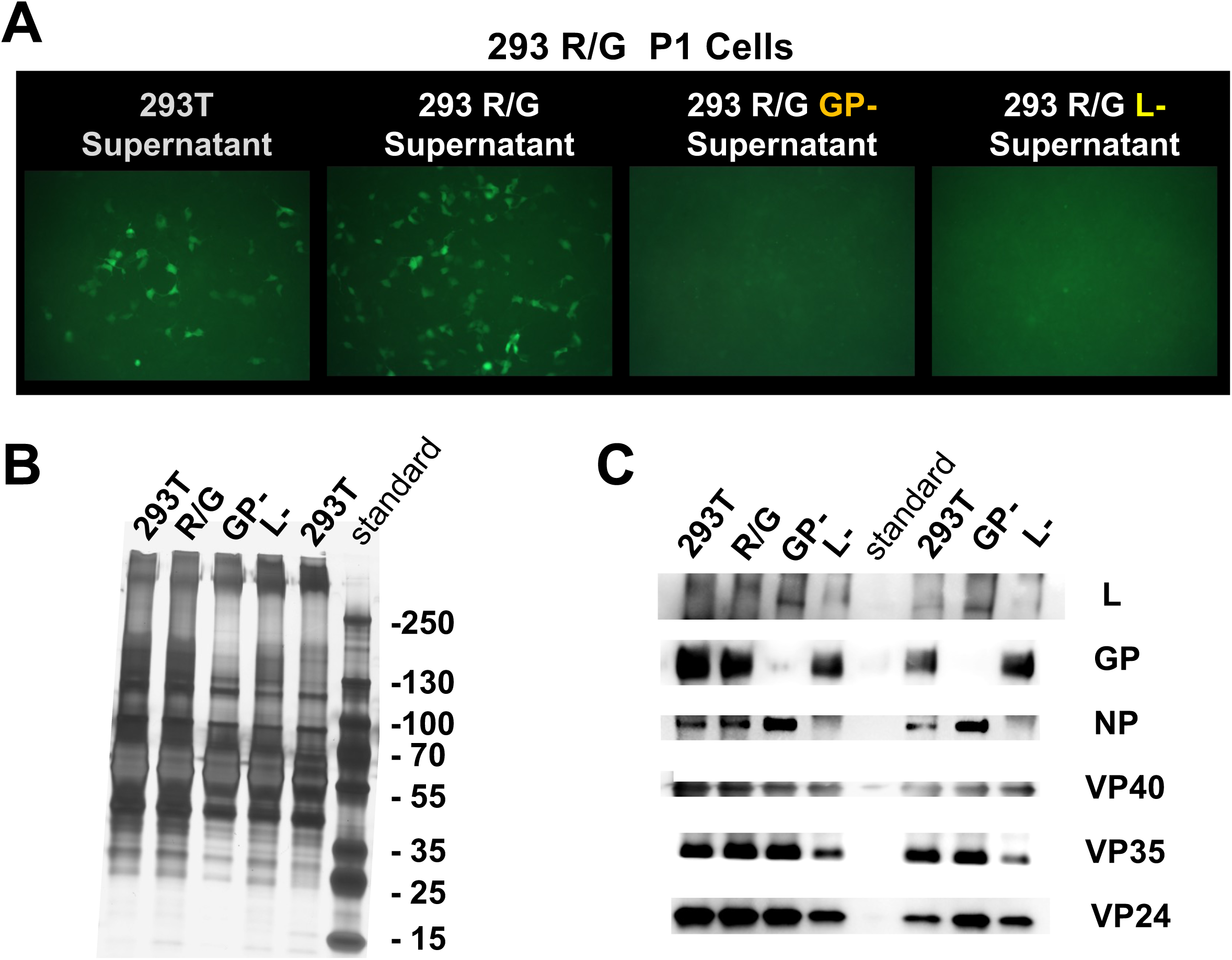
Functional and Protein Characterization of Complete, GP-deficient, and L-deficient EBOV trVLP Preparations. **A)** Representative GFP fluorescence images of P1 R/G reporter cells following exposure to concentrated preparations generated from 293T R/G cells transfected with the complete plasmid set or with plasmid combinations lacking GP (−GP) or L (−L). Concentrated supernatants from 293T and 293T R/G cells are shown as controls. GFP reporter activation was observed following treatment with the complete trVLP preparation but was not detected in the −GP or −L groups. **B)** Silver staining of the corresponding concentrated preparations showing their overall protein profiles. **C)** Western blot analysis of the indicated EBOV proteins of GP, NP, VP40, VP35, VP24, and L in the corresponding preparations.

Approximately 200 µg of each of these concentrated samples was used for intradermal injections into LSL-Luc Cre reporter mice (**Fig. 7**). Under these conditions, concentrated trVLPs produced in 293T or 293 R/G cells produced luciferase activity in the skin with a similar kinetic as was observed previous (**Fig. 3**). In contrast, concentrated trVLPs produced without either GP or without L mediated no detectable luciferase activity.

**Figure 7.**
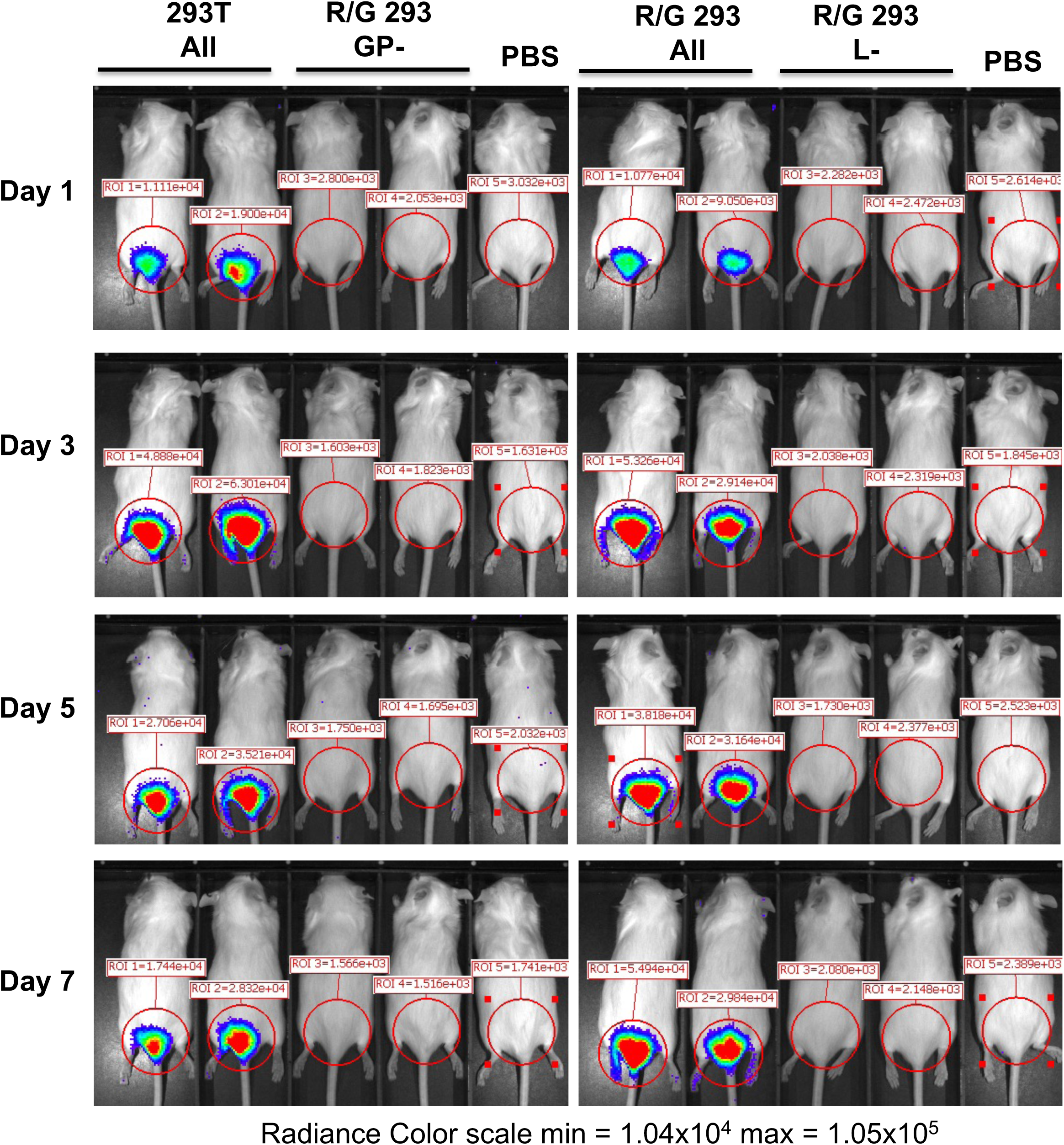
*In vivo* Evaluation of Complete, GP-deficient, and L-deficient EBOV trVLP Preparations. Representative longitudinal bioluminescence images of Cre reporter mice following intradermal administration of complete trVLPs produced in 293T cells, GP-deficient (−GP) preparations produced in R/G 293 cells, complete trVLPs produced in R/G 293 cells, L-deficient (−L) preparations produced in R/G 293 cells, or PBS. Mice were imaged on days 1, 3, 5, and 7 after administration. Red circles indicate the regions of interest (ROIs) used to quantify luciferase activity at the injection site. Two mice were included in each trVLP treatment group, with one PBS control mouse included for each experimental set. Radiance Color scale min = 1.04x10^4^, max = 1.05x10^5^.

### Use of Cre trVLPs to Test Vaccine Efficacy

Finally, we evaluated whether the Cre EBOV trVLP reporter platform could be applied to assess antiviral therapies. In this case, we tested if the system could be used to document vaccine efficacy using a single-cycle adenovirus (SC-Ad) vaccine we developed previously that expresses the 2014 EBOV glycoprotein (SC-Ad-EBOV-GP)[17] (**Fig. 8**).

**Figure 8.**
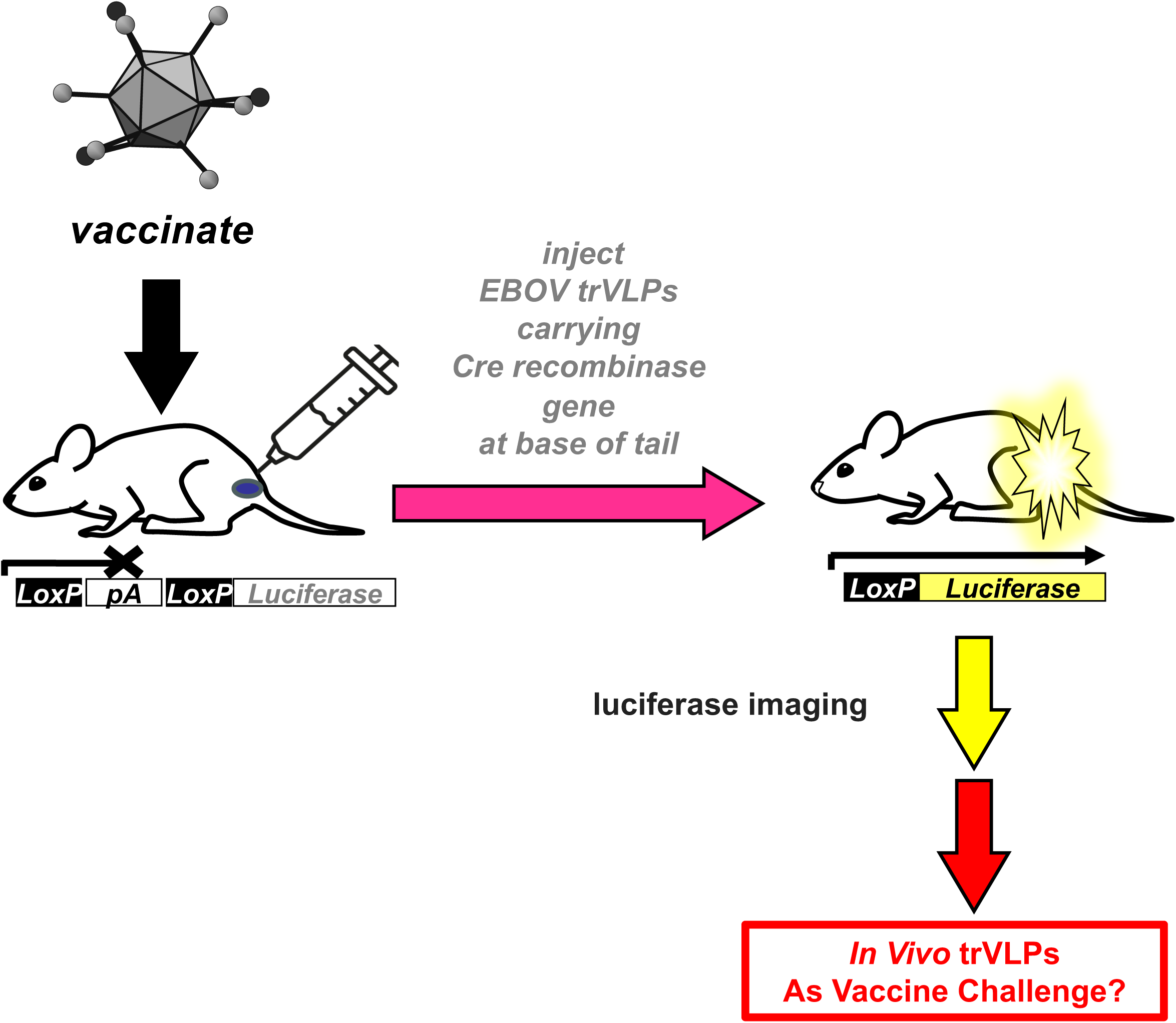
Schematic of the Cre reporter mouse model for evaluating EBOV trVLP challenge *in vivo*. Cre reporter mice were used to assess reporter activation following EBOV trVLP challenge. Delivery of Cre recombinase by trVLPs induces Cre-mediated recombination and activation of luciferase expression in targeted cells. Reporter activation was subsequently monitored by *in vivo* bioluminescence imaging to evaluate trVLP challenge in vaccinated mice.

LSL-Luciferase Cre reporter mice were vaccinated intramuscularly with PBS or with 10^10^ virus particles (vp) of SC-Ad-EBOV-GP. 6 weeks later, at a time when single SC-Ad vaccine immune responses plateau [17], the mice were challenged with concentrated Cre trVLPs and luciferase imaging was performed on subsequent days (**Fig. 9**). Under these conditions, single SC-Ad-EBOV-GP vaccination mediated detectable reductions in luciferase activity in the mice when compared with PBS control vaccine mice. When luciferase activity was compared over time by two-way ANOVA, SC-Ad-EBOV-GP vaccinated mice had significantly lower trVLP activity than PBS controls (p < 0.001), These data suggest that the BSL2 trVLP Cre reporter system can be a potential method to evaluate and screen therapeutics and vaccines against orthoebolaviruses.

**Figure 9.**
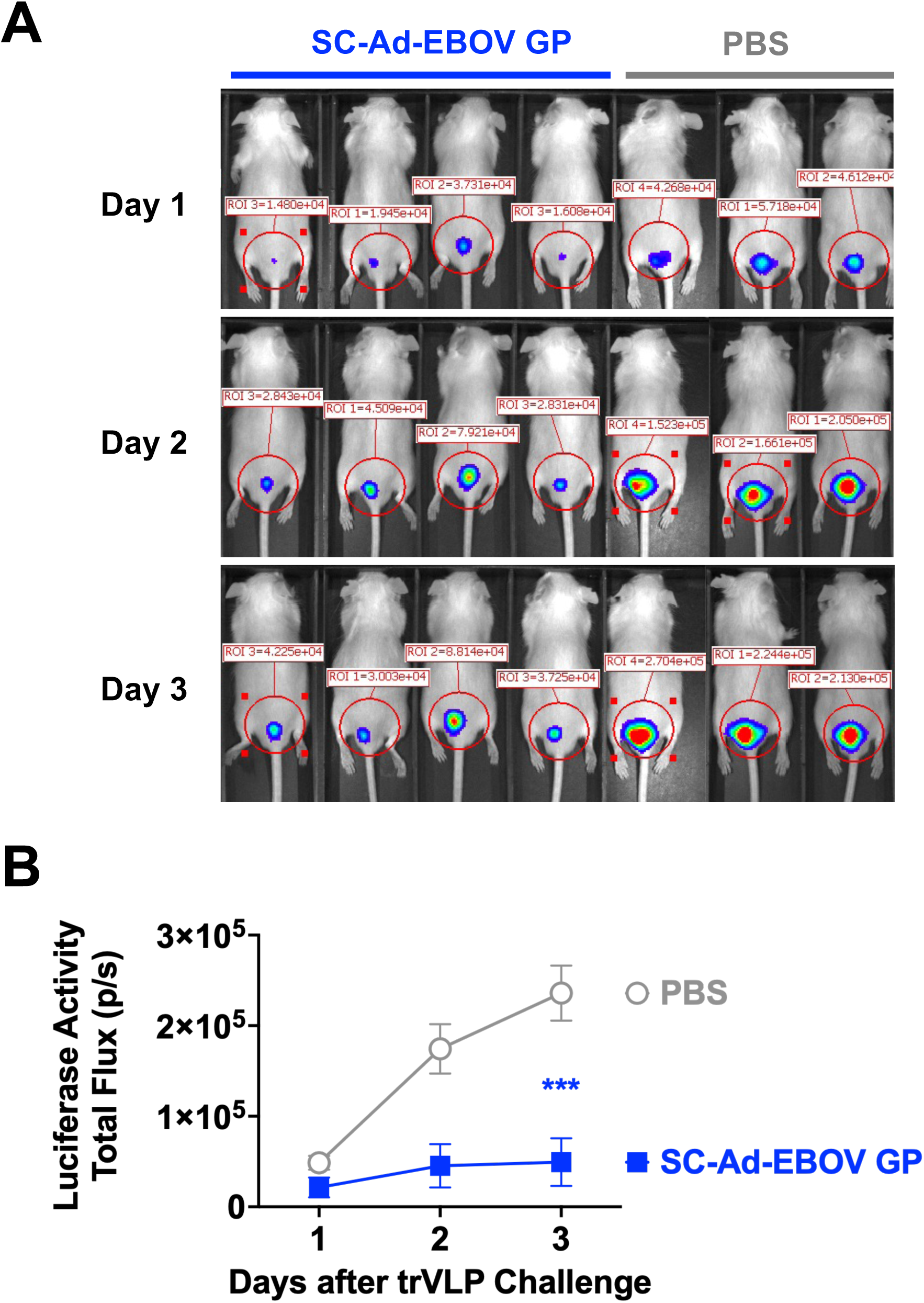
Effects of EBOV Vaccination on Cre trVLPs-induced Luciferase Reporter Activity *in vivo*. LSL-luciferase reporter mice were immunized intramuscularly with SC-Ad6-EBOV GP (10^10^ virus particles [vp]) or PBS. 3 weeks later after immunization, the mice were challenged intradermally with concentrated Cre EBOV trVLPs, and luciferase activity was monitored by longitudinal in vivo bioluminescence imaging on the indicated days. **A)** Representative IVIS images of luciferase activity in SC-Ad6-EBOV GP-immunized and PBS control mice. Red circles indicate the regions of interest (ROIs) used for quantification at the injection site. Radiance Color scale min=1.06e^5^, max=1.10e^6^. **B)** Quantification of whole-animal luciferase activity as total flux (photons/s) over time. Data are presented as mean ± SEM. Statistical comparisons were performed using two-way repeated-measures ANOVA followed by Šídák’s multiple-comparisons test to compare SC-Ad-EBOV GP-immunized mice with PBS controls at each time point. \*\*\**p* < 0.001.

## Discussion

This work was motivated by a desire to test therapeutics and vaccines that could be used against EBOV and BDBV. Most investigators are not part of BSL4 facilities that are needed to test therapeutics against legitimate orthoebolaviruses. Given the danger of working with these BSL4 agents, it is rightly difficult to access these facilities. This low access by most researchers drastically lowers the number of therapeutic or vaccine “hits” that can be screened to identify new or more robust therapies or vaccines. This low access to BSL4 facilities is made even more difficult when there is an ongoing viral outbreak (i.e. currently during the current BDBV outbreak).

Given these impediments and our previous use of Cre reporter systems to track the tropism of RNA and gene therapy vectors [10–14], we tested if Cre-expressing EBOV trVLPs could also be used *in vivo* in Cre reporter mice.

Ebola virus transcription- and replication-competent virus-like particles (trVLPs) have provided an important BSL2-compatible system for investigating multiple stages of the viral life cycle. However, their use has remained predominantly confined to cultured cells and endpoint measurements. This limitation has made it difficult to address several interconnected questions within a single experimental system: how trVLP-associated infection evolves over time in a living host, which immune-cell populations are involved at the site of administration and in draining lymphoid tissues, and how protective interventions alter these processes. The principal advance of the present study is the integration of these previously separate dimensions into a unified *in vivo* reporter framework. By coupling Cre-expressing EBOV trVLPs with Cre-responsive reporter cells and mice, this system enables longitudinal visualization of infection-associated events, identification of infected immune-cell populations, and quantitative assessment of vaccine-induced protection.

A central strength of this framework is that it connects temporal and cellular measurements that are usually obtained independently. Whole-body bioluminescence imaging enables repeated assessment of reporter activity in the same animal and therefore captures the evolution of trVLP-associated infection over time. In contrast, flow cytometry provides cellular resolution but requires terminal tissue collection. Combining these approaches allows macroscopic infection kinetics to be interpreted together with the immune-cell populations identified in the skin and draining lymph nodes. This integration is important because longitudinal imaging alone cannot determine which cells contribute to the observed signal, whereas endpoint immune profiling alone cannot reconstruct the preceding temporal course of infection. The reporter system therefore links organism-level dynamics with cell-level information within the same experimental framework.

The use of irreversible Cre-mediated recombination provides an additional advantage over conventional transient reporter systems. Once Cre is delivered and functional recombination occurs, reporter conversion is permanently recorded in the responsive cell. This genetic memory allows previous infection-associated events to remain detectable even after Cre expression or other transient viral signals have declined. In this context, the reporter does not simply measure the intensity of viral gene expression at a single time point; it records that a cell has received functional Cre delivered through the trVLP system. Such permanent labeling is especially useful for longitudinal studies and for subsequent phenotypic characterization of infected cells by flow cytometry. It also helps explain why the platform can connect early delivery events with later cellular analyses.

Our immune-cell analyses demonstrate that EBOV trVLP-associated reporter conversion can be detected in multiple leukocyte populations within both the skin and draining lymph nodes following intradermal administration. These tissues represent biologically relevant compartments for local antigen capture, innate immune sensing, and initiation of downstream immune responses. The presence of infected reporter-positive cells in the skin is consistent with direct exposure at the administration site, whereas detection in draining lymph nodes suggests involvement of cells within the local lymphatic immune network. Depending on the final distribution observed among dendritic cells, macrophages, Langerhans cells, monocytes, lymphocytes, or other subsets, these findings may provide initial insight into the cellular pathways through which trVLP-associated signals are retained or transported between peripheral tissue and lymphoid organs. The detection of reporter-positive CD207⁺ populations in both the skin and draining lymph nodes is of particular interest given the established role of Langerhans cells and related dendritic-cell populations in antigen capture and communication between peripheral tissues and draining lymphoid organs. In the present study, CD207⁺ cells were analyzed within two phenotypically defined myeloid/DC compartments rather than as functionally distinct Langerhans-cell subsets. The overlapping expression of CD11b, CD11c, MHC-II, and CD207 across cutaneous antigen-presenting cells should therefore be considered when interpreting these populations. Nevertheless, the presence of reporter activation within CD207⁺ compartments illustrates the ability of the platform to interrogate antigen-presenting cell populations that may participate in local trVLP handling and downstream immune responses. Nevertheless, the present study was designed primarily to establish the analytical capability of the platform rather than to define the complete cellular tropism or migration pathway of EBOV. Additional lineage-resolved and spatial studies will be required to distinguish direct infection from cellular trafficking or other mechanisms contributing to reporter-positive populations in draining lymph nodes.

The vaccine experiment further demonstrates that the framework can connect an intervention with both infection-associated and immunological outcomes. Reduced longitudinal reporter signals in vaccinated animals indicate that protective immunity can be detected dynamically rather than inferred only from a terminal endpoint. The accompanying ELISA results provide an independent measure of the vaccine-induced humoral response and strengthen the interpretation that altered reporter kinetics are associated with immunization. Importantly, the value of the platform is not limited to vaccine testing. The same experimental framework could be applied to compare antiviral compounds, neutralizing antibodies, immunomodulatory strategies, or alternative trVLP formulations, provided that the intervention is expected to modify reporter-associated infection events. Longitudinal measurements within the same animal may also reduce inter-animal variability and improve the efficiency of comparative studies.

Several limitations should be considered when interpreting this system. First, reporter conversion is a genetic record of successful functional Cre delivery and recombination rather than a direct quantitative measurement of ongoing viral replication. Reporter-positive cells can therefore reasonably be described as infected within the operational definition of this trVLP reporter system, but reporter intensity should not automatically be interpreted as proportional to active replication at the time of tissue collection. Second, irreversible labeling preserves earlier events and may therefore include cells in which active trVLP-associated gene expression has already ceased. This is an advantage for recording infection history, but it must be considered when interpreting late time points. Third, the immune-cell analysis was limited to the markers and tissues included in the present study. More comprehensive phenotyping, spatial imaging, viral RNA detection, or single-cell transcriptomic analysis would be required to define cellular tropism and migration pathways with greater resolution. Finally, although the system reproduces important features of the EBOV life cycle, it remains a trVLP surrogate and cannot fully model the pathogenicity, systemic dissemination, or disease manifestations of authentic EBOV infection.

These limitations also define several directions in which the framework can be expanded. Incorporating additional Cre-responsive reporter configurations may permit tissue-specific or cell-type-specific analyses. Combining permanent reporter labeling with viral RNA or protein detection could distinguish historical infection events from active viral activity. Spatial imaging of the skin and draining lymph nodes could clarify whether reporter-positive cells arise locally or migrate between compartments. The platform may also facilitate comparisons among trVLP variants, routes of administration, host backgrounds, and candidate countermeasures. Although the present study focuses on EBOV, the underlying strategy may be adaptable to other engineered viral surrogate systems in which functional delivery of Cre can be coupled to a responsive host reporter.

In conclusion, this study extends EBOV trVLP research beyond static measurements in cultured cells and establishes a BSL2-compatible framework for dynamic investigation in living animals. By integrating longitudinal imaging, permanent genetic recording, immune-cell profiling, and intervention assessment, the system enables infection dynamics, host-cell involvement, and protective responses to be examined as connected components of the same biological process. More broadly, this work shifts EBOV trVLP studies from isolated measurements of viral activity toward integrated analysis of virus–host–Most importantly, these questions can be investigated under BSL2 conditions to allow more researchers to test their leads, but also to reduce the burden of hits needing to be tested against orthoebolaviruses by already over-burdened scientists at BSL4 facilities.

## Materials and Methods

### Plasmids

EBOV (Mayinga variant) protein-expressing plasmids pCAGGS-NP, -VP35, -VP30, and-L were previously generated and described [9, 18]. To compliment these, VP24 and VP40 expression plasmids were constructed and used for co-transfection. Plasmid pATX-T7-MG-Cre was constructed for T7 polymerase-driven expression of a negative stranded monocistronic Cre expressing mini minigenome (1×MG-Cre). Cre reporter plasmids is a lentiviral vector carrying EF1a-loxP-DsRedExpress2-loxP-EGFP Cre Reporter that was a gift from Niels Geijsen (Addgene plasmid # 62732; http://n2t.net/addgene:62732; RRID:Addgene_62732)[19]. These plasmids were prepared as endotoxin-free DNAs and the plasmid sequences were confirmed via Sanger sequencing or next-generation sequencing before use.

### Cell culture

293 Red/Green Cre reporter cells (R/G) cells were produced by infection of 293 cells with Cre reporter lentivirus that carries EF1a-loxP-DsRedExpress2-loxP-EGFP. When these cells are infected with a Cre-expressing vector, the recombinase excises the loxP-flanked red fluorescent protein (RFP) STOP cassette and switches fluorescence from red to green fluorescent protein (GFP) expression. 293 R/G and 293T cells were maintained in Dulbecco’s Modified Eagle Medium (Gibco, Waltham, MA, USA) supplemented with 10% (v/v) fetal bovine serum and 1% (v/v) penicillin and streptomycin. Cells were incubated in 5% CO2 at 37°C.

### Cre-expressing EBOV trVLP production and purification

For P0 transfections, 293T or 293 R/G cells were seeded in 6-well plates. After 24 hours, their media was changed to 2% FBS DMEM and the cells were transfected with a 3:1 ratio of TransIT-LT1 Transfection Reagent (Mirus Bio, Madison, WI, USA) with plasmid DNA in OptiMEM media (Gibco, Waltham, MA, USA). Plasmid amounts each well for the plasmids were: pCAGGS-NP 250 ng, pCAGGS-VP35 250 ng, pCAGGS-VP30 150 ng, pCAGGS-L 2000 ng, pCAGGS-GP 500 ng, pCAGGS-VP40 500 ng, pCAGGS-VP24 15 ng and 1×MG-Cre 250 ng. For experiments utilizing a T7 polymerase-driven minigenome, 500 ng of pCAGGS-T7pol was included in transfection reactions. Their media was changed after 8 hours and in some cases GFP and RFP expression was monitored by fluorescence microscopy. After 72-120 hours, cell supernatants were harvested and clarified by centrifugation at 1000 g for 10 minutes. For concentrated trVLPs, 7 ml of clarified trVLP supernatants were applied to a 3 ml 20% sucrose layer in (-/-) PBS buffer (calcium- and magnesium-free phosphate-buffered saline) in a Beckman centrifuge tube (Cat. No. C14293) and centrifuged at 32,000 rpm in a SW32-TI rotor for 2 hours at 4°C using a Beckman L7 ultracentrifuge. Pellets were resuspended in 300 µL of (-/-) PBS buffer, then injected in mice or treated in P1 cells. The rest trVLP preparations were aliquoted and stored at −80 °C.

Total protein concentrations were determined using BCA assay (Pierce BCA Protein Assay Kits, Thermo Fisher Scientific, cat. no. 23225). Silver staining (Pierce Silver Stain Kit, Thermo Fisher Scientific, cat. no. 24612) and western blot were performed to assess individual viral proteins.

### trVLP infections

40 µL resuspended trVLPs prepared in the previous description were transferred to wells of 293 R/G cells (P1 cells) in 6-well plate and incubated at 37°C for the indicated times.

### Cre reporter mice

mT/mG mice (aka. B6.129(Cg)-Gt(ROSA)26Sortm4(ACTB-tdTomato,-EGFP)Luo/J strain # 007676) and LSL-Luc mice (aka FVB.129S6(B6)-Gt(ROSA)26Sortm1(Luc)Kael/J mice strain # 005125) were purchased from The Jackson Laboratory (Bar Harbor, ME, USA). mT/mG, and the LSL-Luc mice were crossed as F1 progeny for the experiments involving combined luciferase imaging and flow cytometry. Mice were housed in the Mayo Clinic Animal Facility. All animal handling and experiments were carried out according to the provisions of the Animal Welfare Act, PHS Animal Welfare Policy, the principles of the NIH Guide for the Care and Use of Laboratory Animals, and the policies and procedures of the Institutional Animal Care and Use Committee at Mayo Clinic.

### *In vivo* reporter mouse experiments

Cre reporter mice were injected intradermally at the base of the tail with 100 µL of the indicated trVLP preparation (∼200 µg total protein, as determined by BCA assay) or PBS. Luciferase expression was assessed by bioluminescence imaging on days 1, 3, 5, and 7 after injection, as previously described [14]. Briefly, mice were anesthetized with isoflurane and administered D-luciferin potassium salt (150 mg/kg body weight in PBS; GoldBio) by intraperitoneal injection. After 15 min, dorsal images were acquired using a Lumina S5 IVIS Spectrum system (Revvity) under standardized imaging parameters. On day 7, immediately after whole-body imaging, mice were euthanized, and the abdominal skin was opened to expose the inguinal lymph nodes, which were imaged using the same instrument settings. Bioluminescent signals were quantified using Living Image software (v.4.7.3, Revvity) with consistent regions of interest (ROIs) applied across samples. Following imaging, skin from the injection site and draining inguinal lymph nodes were collected and processed for flow cytometric analysis.

### Skin and lymph node immune-cell analysis

Skin and inguinal lymph nodes were processed into single-cell suspensions as previously described [14]. Briefly, skin tissue was dissociated using the Epidermis Dissociation Kit (mouse; Miltenyi Biotec), and cells from skin and lymph nodes were washed and resuspended in staining buffer (PBS containing 2% FBS). Cell viability was assessed using Ghost Violet 510 Viability Dye (Tonbo Biosciences), followed by surface staining for 30 min at 4°C in the dark. The antibody panel included anti-mouse CD45 (FITC), CD3 (BV510), CD11b (PE), CD11c (PerCP), MHC-II (BV421), and CD207/Langerin (APC) (BioLegend, San Diego, CA, USA). After staining, cells were fixed with 2% paraformaldehyde for 30 min on ice, washed, and resuspended for flow cytometric analysis. Data were acquired using a Cytek Northern Lights spectral flow cytometer and analyzed using FlowJo software. Spectral unmixing was performed using appropriate single-stained reference controls. Doublets were excluded by FSC-A versus FSC-H gating, and dead cells were excluded based on Ghost Violet 510 staining. CD45 expression was used to distinguish immune and non-immune cell populations, and immune-cell subsets were further analyzed based on the expression of CD3, CD11b, CD11c, MHC-II, and CD207. GFP expression was subsequently assessed within the indicated cell populations. GFP-positive gates were established using PBS-injected control mice to define background fluorescence and were applied consistently across all experimental samples. Fluorescence-minus-one controls were included for CD207. Experiments were independently repeated three times.

### EBOV vaccine construction and valuation

Codon-optimized glycoprotein (GP) cDNA encoding EBOV-Makona isolate KJ660346 [20] (Genscript) was recombined into single-cycle adenovirus serotype 6 (SC-Ad6) to generate SC-Ad6-EBOV-GP as previously described in [17].

GP expression was confirmed by Western blot analysis of SC-Ad6-EBOV-GP-infected A549 cells. Mice were immunized intramuscularly with e^10^ vp/mice SC-Ad6-EBOV-GP 3 weeks before subsequent experimental procedures. Luciferase expression was assessed via bioluminescence imaging at day 1, 2, 3 after trVLPs injection.

### Western blot

Samples were mixed with 4× Laemmli sample buffer (Bio-Rad, Hercules, CA, USA) containing 10 mM DTT and heated for 10 min before separation by SDS-PAGE. Proteins were transferred onto PVDF membranes using a semi-dry transfer system at 15 V for 32 min. Membranes were blocked with 5% (w/v) skim milk in TBS-Tween and incubated overnight at 4°C with anti-VP40 (clone 5B12; IBT BioServices; Cat. No. 0201-017), anti-VP24 (polyclonal; Sino Biological; Cat. No. 40454-T46), anti-NP (custom rabbit polyclonal antibody against the NP peptide Cys-PAVSSGKNIKRT; Biomatik, Kitchener, ON, Canada), anti-EBOV GP (rabbit polyclonal; IBT BioServices, Rockville, MD, USA), anti-VP35 (rabbit polyclonal; IBT BioServices; Cat. No. 0301-040), and anti-EBOV L polymerase (rabbit polyclonal; IBT BioServices; Cat. No. 0301-045) primary antibodies. Membranes were washed three times with TBS-Tween for 10 min each and subsequently incubated with appropriate HRP-conjugated secondary antibodies. Protein signals were detected using SuperSignal West Femto chemiluminescent substrate (Thermo Fisher Scientific, Waltham, MA, USA) and imaged using a Bio-Rad ChemiDoc imaging system.

### Statistical analysis

For GFP fluorescence intensity in P0 reporter cells, data are presented as mean ± SD from three independent experiments. Statistical comparisons were performed using ordinary one-way ANOVA followed by Dunnett’s multiple-comparisons test for P0 cells, with each time point compared with the PBS control group. For P1 cells, data are presented as mean ± SD, and statistical comparisons were performed using a two-tailed unpaired t-test with Welch’s correction. For luciferase activity, data are presented on the original scale as mean ± SEM. Longitudinal data with missing values were analyzed using a repeated-measures mixed-effects model (REML), followed by Tukey’s multiple-comparisons test. For complete longitudinal datasets, two-way repeated-measures ANOVA followed by Šídák’s multiple-comparisons test was used to compare treatment groups at each time point. The flow cytometry data are presented as percentages of the indicated cell populations. For comparisons among multiple groups, the nonparametric Kruskal–Wallis test followed by Dunn’s multiple comparisons test was applied, with all pairwise group comparisons performed. The data are presented as individual points representing each mouse, with mean ± SD. Statistical analysis for this study was performed using GraphPad Prism version 10.4.1.3. A *p*-value < 0.05 was considered statistically significant.

## Acknowledgements

This work was supported by grant NIH/NIAID R21 AI188373 to M.A.B. We would like to acknowledge the excellent technical assistance of Mary E. Barry and Zoé Dubin.

## Author contributions

Conception and Design: J.L, B.N.Z., and M.A.B. Development of Methodology: J.L, B.N.Z., and M.A.B. Acquisition of Data: J.L Analysis and Interpretation of Data: J.L, B.N.Z., L.W., and M.A.B. Writing, Review, and/or Revision of the Manuscript: J.L, B.N.Z., L.W., and M.A.B. Artistic renderings: J.L. Study Supervision: M.A.B.

---

**Supplemental Figure 1.**
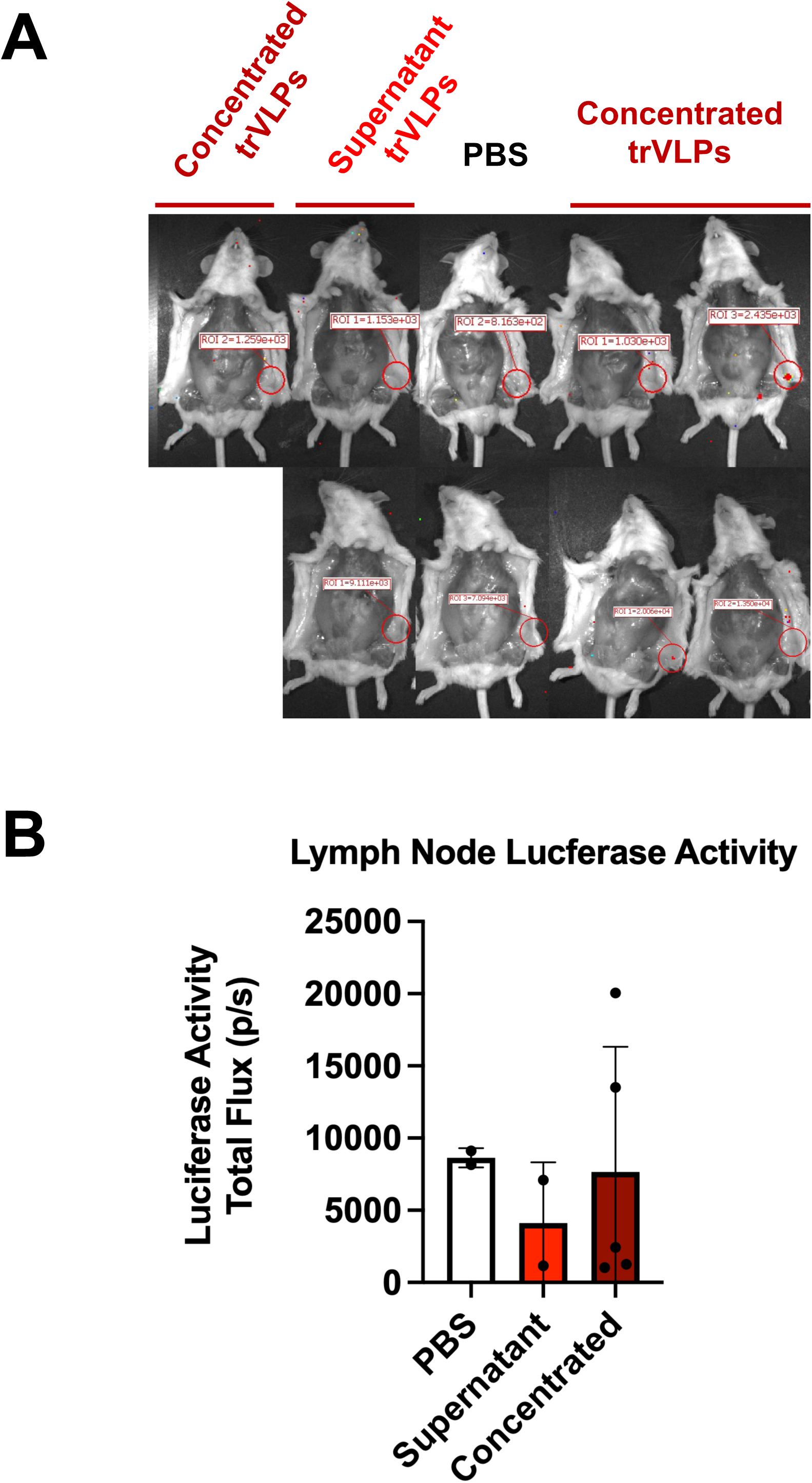
Luciferase Activity in Skin-draining Inguinal Lymph Nodes. **A)** Representative bioluminescence images of the draining inguinal lymph nodes on day 7. Following euthanasia, the abdominal skin was opened to expose the inguinal lymph nodes, which were imaged using IVIS. Red circles indicate the regions of interest (ROIs) used for quantification of luciferase activity. **B)** Quantification of luciferase activity in the draining inguinal lymph nodes. Detectable luciferase signals were observed in two individual mice, with no statistically significant differences among treatment groups. One mouse in the concentrated trVLP group died on day 5 and was therefore not included in the day 7 lymph-node imaging.

**Supplemental Figure 2.**
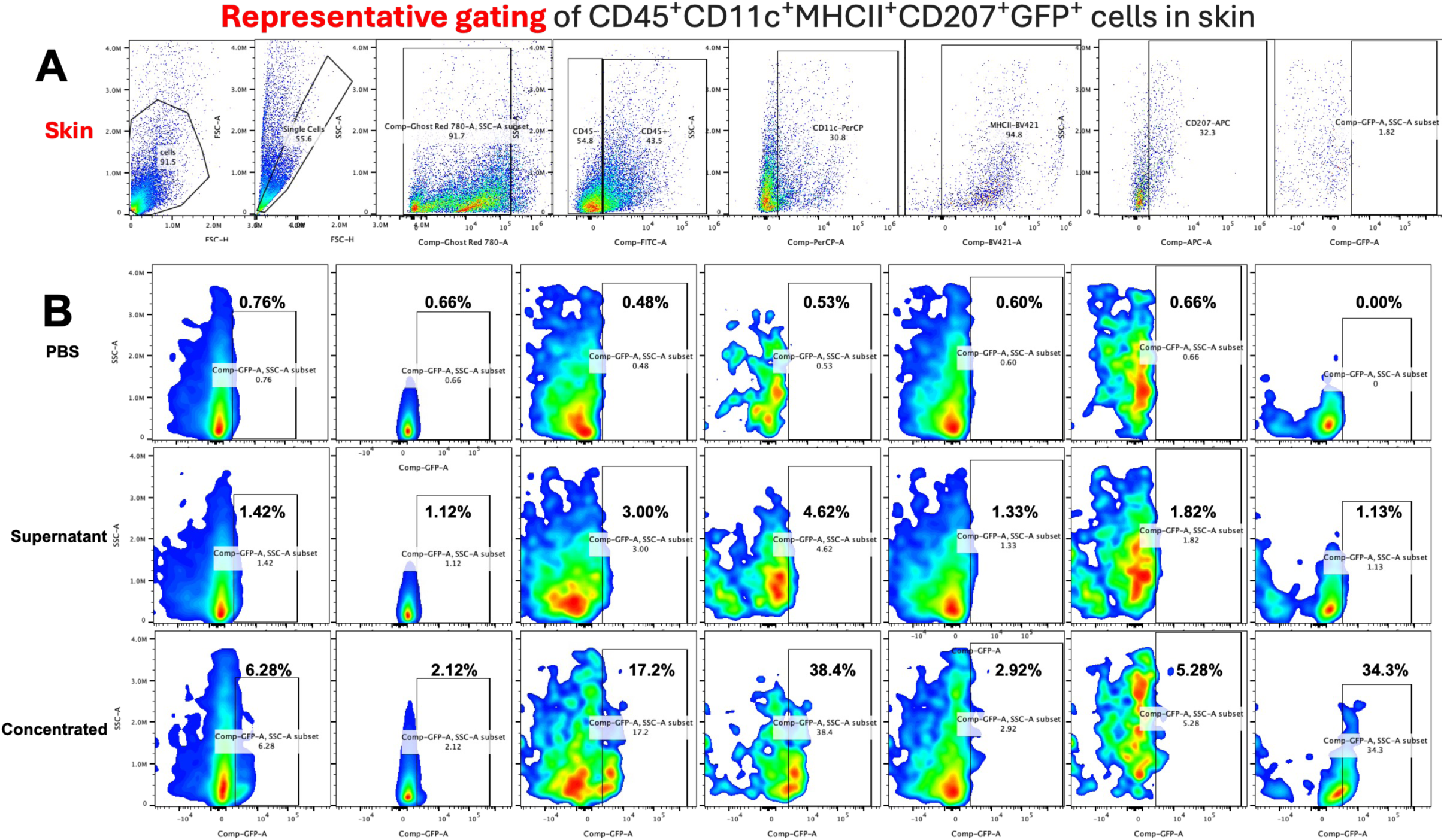
Representative Flow Cytometry Gating for Skin Samples. Representative gating strategy for analysis of GFP-positive immune-cell populations in skin. Following exclusion of debris, doublets, and nonviable cells, CD45⁺ immune cells were identified and sequentially gated based on CD11c, MHC-II, CD207, and GFP expression. Representative GFP plots from mice receiving PBS, unconcentrated trVLP-containing supernatant, or concentrated trVLPs are shown.

**Supplemental Figure 3.**
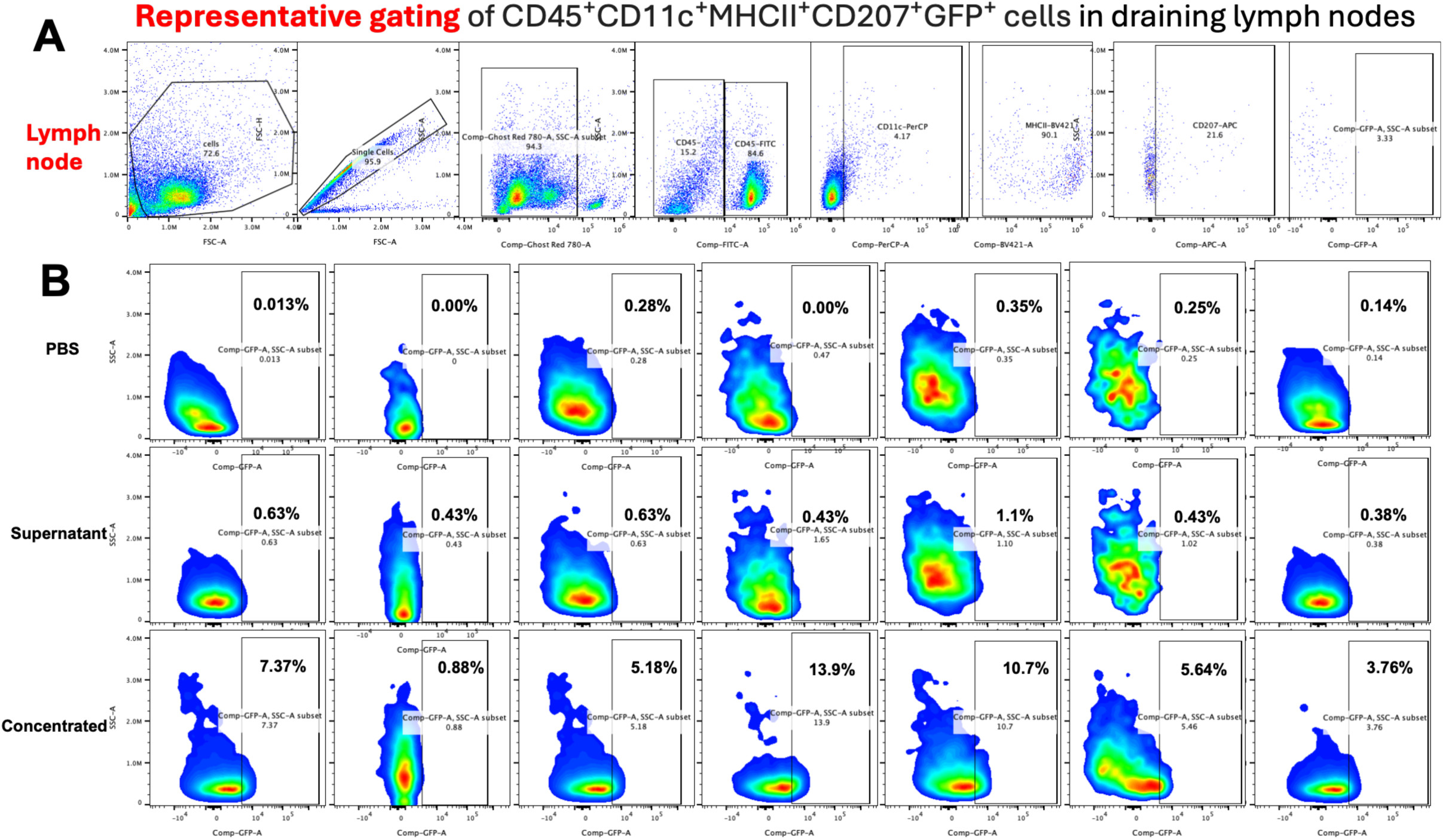
Representative Flow Cytometry Gating for Draining Lymph Node Samples. Representative flow cytometry plots of draining inguinal lymph node cells from mice receiving PBS, unconcentrated trVLP-containing supernatant, or concentrated trVLPs. The gating strategy was performed as described in Supplemental Figure 2.

**Supplemental Figure 4.**
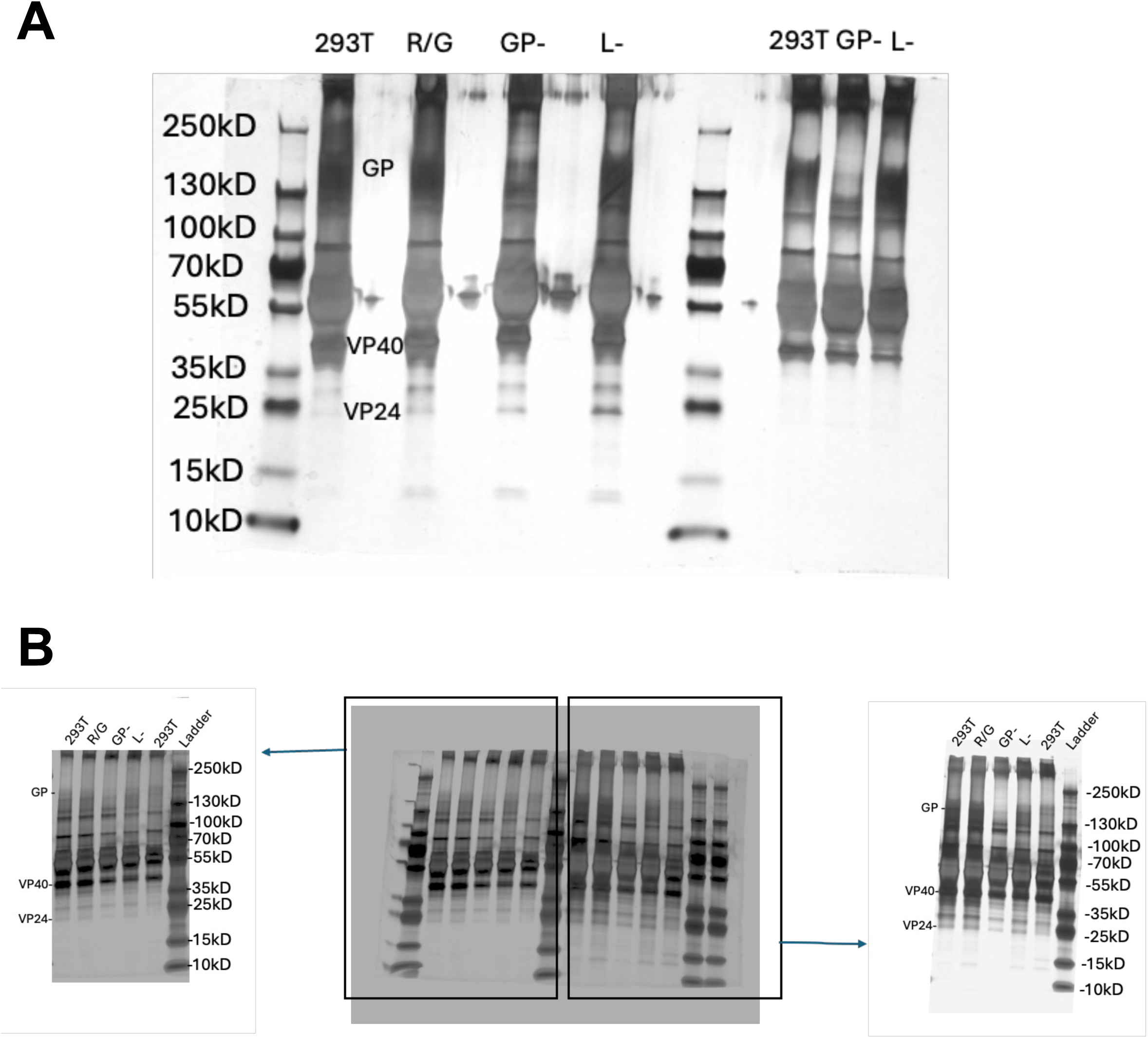
Original silver-stained SDS-PAGE gels of EBOV trVLP preparations. Original and representative silver-stained SDS-PAGE gel images used for characterization of the indicated trVLP preparations. Molecular-weight markers are shown in kilodaltons (kDa). The approximate migration positions of GP, VP40, and VP24 are indicated. The uncropped gel images are provided to show the overall protein profiles of the analyzed preparations.

**Supplemental Figure 5.**
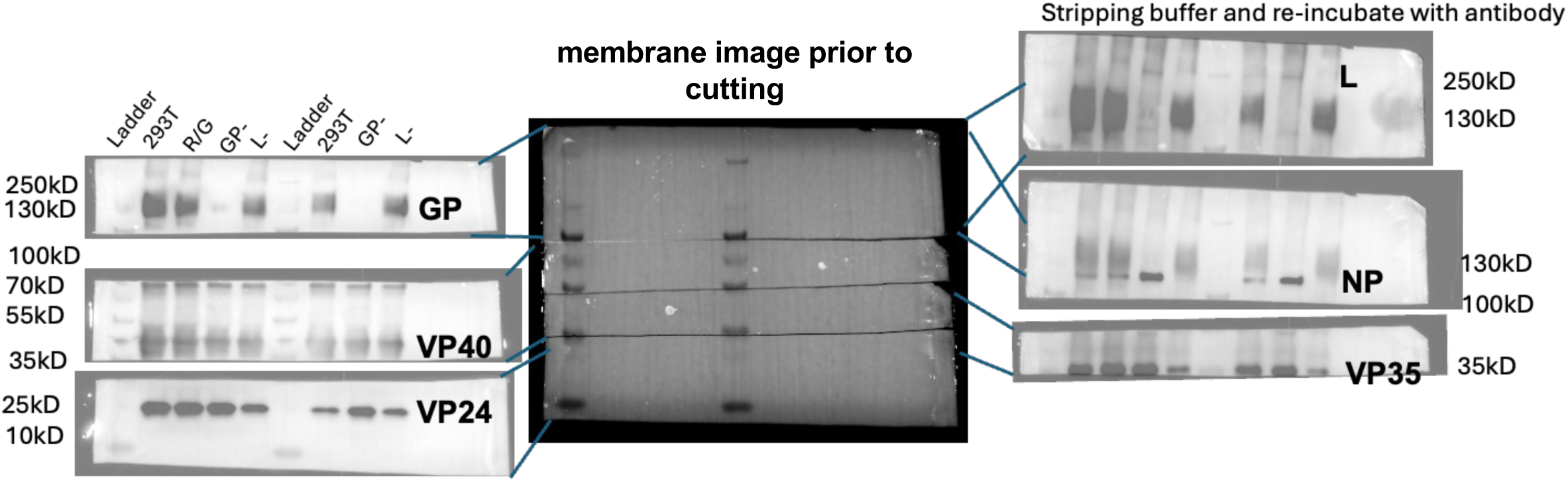
Original Western blot images of EBOV trVLP proteins. Original membrane and Western blot images used for characterization of the indicated trVLP preparations. The membrane image prior to sectioning and the corresponding immunoblots probed for individual EBOV proteins are shown. The positions of GP, VP40, VP24, L, NP, and VP35 are indicated.

